# tACS phase-specifically biases brightness perception of flickering light

**DOI:** 10.1101/2021.06.26.450021

**Authors:** Marina Fiene, Jan-Ole Radecke, Jonas Misselhorn, Malte Sengelmann, Christoph S. Herrmann, Till R. Schneider, Bettina C. Schwab, Andreas K. Engel

**Author notes:** Department of Psychiatry and Psychotherapy, University of Lübeck, Lübeck, 23562 Germany (Jan-Ole Radecke). Biomedical Signals and Systems, Technical Medical Center, University of Twente, Enschede, 7522 The Netherlands (Bettina C. Schwab). shared last authorship (Bettina C. Schwab, Andreas K. Engel). Corresponding author: Marina Fiene.

## Abstract

**Background:** Visual phenomena like brightness illusions impressively demonstrate the highly constructive nature of perception. In addition to physical illumination, the subjective experience of brightness is related to temporal neural dynamics in visual cortex.

**Objective:** Here, we asked whether biasing the temporal pattern of neural excitability in visual cortex by transcranial alternating current stimulation (tACS) modulates brightness perception of concurrent rhythmic visual stimuli.

**Methods:** Participants performed a brightness discrimination task of two flickering lights, one of which was targeted by same-frequency electrical stimulation at varying phase shifts. tACS was applied with an occipital and a periorbital active control montage, based on simulations of electrical currents using finite-element head models.

**Results:** Experimental results reveal that flicker brightness perception is modulated dependent on the phase shift between sensory and electrical stimulation, solely under occipital tACS. Phase-specific modulatory effects by tACS were dependent on flicker-evoked neural phase stability at the tACS-targeted frequency, recorded prior to electrical stimulation. The optimal timing of tACS application leading to enhanced brightness perception was further correlated with the neural phase delay of the cortical flicker response.

**Conclusions:** Our results corroborate the role of temporally coordinated neural activity in visual cortex for brightness perception of rhythmic visual input in humans. Phase-specific behavioral modulations by tACS emphasize its efficacy to transfer perceptually relevant temporal information to the cortex. These findings provide an important step towards understanding the basis of visual perception and further confirm electrical stimulation as a tool for advancing controlled modulations of neural activity and related behavior.

## Introduction

Dissociations between human brightness estimation and objective viewing conditions reveal important insight into the constructive mechanisms of perceptual processing. Depending on constituents of the visual scene, feedforward projections along the visual pathway are subject to complex neural computations that shape perception in addition to the physical level of illumination [1–4]. A popular approach to examine the neural substrate of perception builds on frequency tagging of neural activity by sinusoidally modulated luminance stimuli that evoke phase-locked responses in visual cortex [5]. This way, singlecell firing patterns were shown to follow the frequency of flickering visual stimuli, reflecting actual as well as illusory brightness percepts [1,6–8]. Correlations between firing pattern modulation and the subjective experience of brightness were found in primary visual cortex but not earlier in the visual pathway. Further, invasive recordings in cats revealed a relation between brightness perception and enhanced discharge rates or synchronization of neural firing at the tagging-frequency in visual cortex [9].

Neural synchronization has long been discussed as a candidate mechanism in the cerebral cortex that increases the efficacy of neural responses in driving target neurons, thereby enhancing response saliency [10–12]. Noninvasive recordings in humans of flicker entrained activity, i.e., visually evoked steady-state responses (SSRs), converge with these findings. Perceptual competition between two concurrently presented flickering stimuli of equal luminance was biased in favor of the stimulus with greater inter-trial phase coherence of evoked SSRs [13,14]. Further, flicker evoked response amplitudes were shown to correlate not only with increases in physical luminance contrast [15–18], but also with the strength of illusory brightness percepts as early as in primary visual cortex [2,3,19,20]. Thus, temporal neural dynamics at early stages of cortical processing are assumed not merely to reflect physical stimulation properties but rather to code the subjective quality of perception. Given this correlative evidence, does a manipulation of flicker-evoked rhythms affect the subjective experience of brightness under otherwise constant viewing conditions?

Transcranial alternating current stimulation (tACS) is used with the aim to modulate neural dynamics by phase-specific excitability modulation in targeted brain regions [21,22]. Invasive recordings in monkeys yielded key evidence that neuronal spike timing follows the phase of the externally applied electric current, thereby inducing higher synchrony and related power increases at the stimulation frequency [23–25]. Only recently, we have shown that pairing tACS with same-frequency flicker results in systematic enhancement and suppression of flicker-evoked SSR amplitudes dependent on the phase shift between visual and electrical stimulation [26]. This finding corroborated the capability of tACS to shape population level neural responses to natural visual inputs in humans [27,28]. Yet, crucially, whether this tACS-induced modulation of neural activity to flickering light also affects the subjective experience of flicker brightness has not been established. Evidence for perceptual deterioration or improvement by interfering with flicker-evoked responses would provide a strong claim about the functional relevance of the underlying neural response.

Here, we asked whether biasing cortical processing of rhythmic visual stimulation by tACS under constant luminance conditions translates into perceptual changes. By using oscillatory entrainment via tACS, we aimed to modulate flicker-evoked responses by concurrent electrically-induced temporal changes in neuronal excitability within the visual cortex. Participants performed a two-flicker brightness discrimination task, while tACS was simultaneously applied at one of the two flicker frequencies with systematic phase shift. We hypothesized that the modulation of neural response dynamics in visual cortex by phase-shifted visually and electrically driven neuromodulation leads to perceptual changes of flicker brightness.

## Materials and Methods

### Participants

38 right-handed participants (mean age 25.74 ± standard deviation 4.06 years, 22 female, 16 male) volunteered for this study. All participants had normal or corrected-to-normal vision and no history of psychiatric or neurological illness. Handedness was assessed via the short version of the Edinburgh Handedness Inventory. Participants gave written informed consent prior to participation and were monetarily compensated. The study was approved by the ethics committee of the Hamburg Medical Association (PV4908) and conducted in accordance with the declaration of Helsinki.

### Experimental design

The experimental protocol was conducted on two separate days for occipital and periorbital tACS in counterbalanced order. In the beginning of each testing session, EEG was recorded in order to assess the physiological response to visual flicker stimulation and the spectral properties of brain activity during resting-state. After psychophysical estimation of brightness discrimination thresholds for individualization of physical luminance intensity (see Supplementary Material A), participants performed the main tACS experiment.

During the tACS experiment, participants were presented with a two-flicker brightness discrimination task between an 8 Hz tACS-targeted flicker (LED_8Hz_) and a 12 Hz reference flicker (LED_12Hz_). Multi-electrode tACS at 8 Hz was simultaneously applied at six different phase shifts (0°, 60°, 120°, 180°, 240°, 300°) relative to the 8 Hz flicker cycle (Fig. 1A). Single flicker trials were presented at five luminance ratios at [35, 42.5, 50, 57.5, 65] % of the individual discrimination performance. The flicker phase angle at light onset of the LED_8Hz_ was set to 0°, while the LED_12Hz_ started randomly at four different phase angles (0°, 90°, 180°, 270°). Single flicker trials had a duration of 2 sec, followed by a break of 1.75 - 2 sec.

**Fig. 1.**
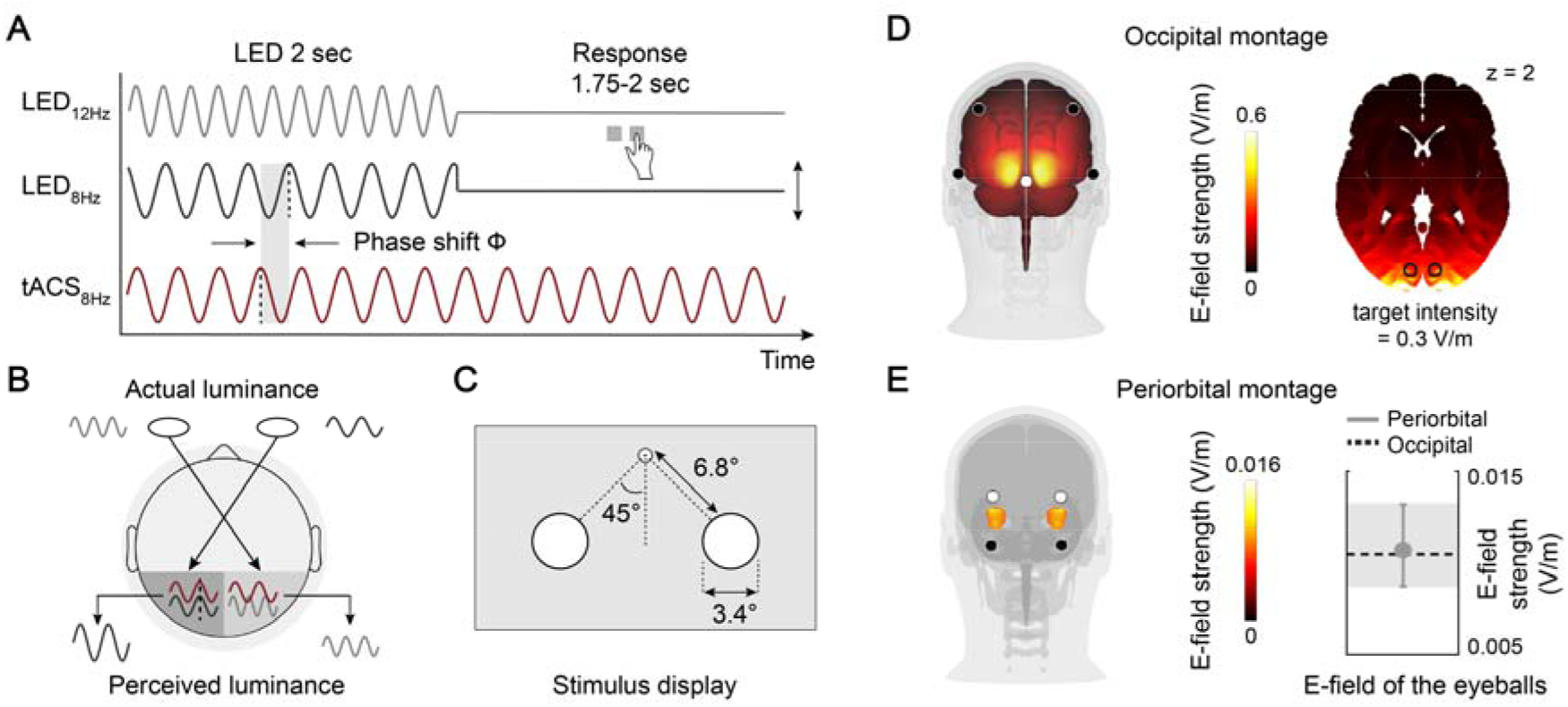
Experimental setup. (A) During continuous transcranial alternating current stimulation (tACS) at 8 Hz, participants were asked to judge per button press which of the two simultaneously presented flickers appeared brighter. Luminance of the LED_8Hz_ was varied while the LED_12Hz_ remained at constant luminance. The lag between LED_8Hz_ and tACS onset was systematically varied across six different phase shifts. (B) Schematic of the experimental hypothesis that phase-specific electrical stimulation of same-frequency flicker-evoked responses systematically modulates brightness perception. (C) The two flickers were presented simultaneously in the lower visual field while participants fixated the upper fixation dot. (D) During occipital tACS at 4 mA peak-to-peak, estimated maximum electric field strength was about 0.6 V/m, reaching 0.3 V/m in the target regions within the upper bank of the calcarine sulcus (MNI-coordinates: −10, −90, 2 and 10, −90, 2). (E) During the periorbital control condition, estimated electric field strength in the eyeballs for tACS intensities at 80% phosphene threshold was comparable to the field magnitude induced by occipital tACS due to current shunting across the scalp (periorbital: bootstrapped mean .01,95%-CI).

After each trial, participants indicated the side of flicker presentation appearing as brighter per button press (or darker, depending on the assigned counterbalanced condition). 19 trials were presented for each of the six tACS-LED_8Hz_ phase shift conditions and five flicker luminance ratios, resulting in 570 trials in total. Trial presentation was subdivided in three blocks à 13.3 min, interrupted by 10 min breaks. Side of flicker presentation and phase offsets were counterbalanced across tACS-LED_8Hz_ phase shift conditions.

### Transcranial electrical stimulation

During the tACS experiment, multi-electrode tACS was applied at 8 Hz via Ag/AgCl electrodes (12 mm diameter) using neuroConn stimulators (DC-Stimulator plus, neuroConn, Illmenau, Germany). One hour before tACS application, the skin beneath the stimulation electrodes was prepared with EMLA cream for local anesthesia to reduce transcutaneous stimulation effects by tACS (2.5 % lidocaine, 2.5 % prilocaine). Electrode impedances were kept below 10 kΩ and comparable between electrodes to ensure an evenly distributed electric field. During the tACS experiment of both sessions, stimulation was applied for 40 min in total, divided in three blocks of 13.3 min duration including a linear ramp-up time of 6 sec and another 6 sec at the final intensity before the start of visual stimulation.

For occipital tACS, current was applied via a 4 x 1 montage over the parieto-occipital cortex at 4 mA peak-to-peak (Fig. 1D). The stimulation montage was tailored to target the upper bank of the calcarine sulcus, as described in detail in Supplementary Material A. Estimated electric field strengths by simulations based on finite-element head models were comparable to electric field strengths shown to successfully modulate neural activity in nonhuman primates and humans [25,29,30]. The periorbital tACS montage served as an active control condition for sub-threshold stimulation effects of the retinae due to current shunting across the scalp (Fig. 1E). Current was applied via four electrodes placed superior and inferior to the eyes with current flow in the vertical direction [31,32]. Stimulation intensities were adjusted to 80% of the individual phosphene threshold [33]. Mean stimulation intensity was 0.071 ± 0.037 mA peak-to-peak. Peripheral tACS side-effects were systematically quantified at the end of each session via a questionnaire, assessing the perceived strength and temporal course of skin sensations, fatigue and phosphene perception (Supplementary Material B, Figure B1). Out of 38 participants, two reported supra-threshold phosphene perception under occipital tACS and were therefore excluded from data analysis.

### Electrophysiological recording and data analysis

EEG data were recorded from 18 Ag/AgCl electrodes (12mm diameter) mounted in parieto-occipital regions in an elastic cap for 64 electrodes (Easycap, Herrsching, Germany). We assessed 60 sec resting state EEG with eyes open and the electrophysiological response to flicker stimulation while participants were presented with an LED flickering at either 8 Hz or 12 Hz for 60 sec in the right lower visual field. Electrode impedances were kept below 20 kΩ. The electrooculogram was recorded with two electrodes placed below the eyes. EEG signals were referenced to the nose tip and recorded using BrainAmp MR plus amplifiers (Brain Products GmbH, Gilching, Germany) and the corresponding software (Brain Products GmbH, Recorder 1.20). Data were recorded with an online passband of 0.016 - 250 Hz and digitized with a sampling rate of 5 kHz.

EEG data were preprocessed and analyzed in Matlab (The MathWorks Inc.) using the analysis toolbox FieldTrip [34] and custom-made scripts. Data were bandpass-filtered between 1 and 20 Hz using a Hamming-windowed sinc FIR (finite impulse response) filter (order 8250) and segmented in 2 sec time epochs. We quantified the extent of visual entrainment by the phase locking value (PLV) between EEG and visual flicker [35]. A Hilbert transform was computed on the flicker and the EEG signal of each epoch and at each electrode. The time-dependent differences of the flicker and EEG signal phase (*ϕ_flicker_t_* – *ϕ_EEG_t_*) for each sample *t* served to compute the PLV

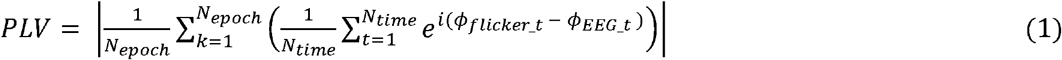

with *N_time_* time steps per epoch, and *N_epoch_* epochs. Comparability of flicker entrainment strength between the two tACS sessions and flicker frequencies was computed by repeated measures analysis of variance (ANOVA).

### Statistical data analysis of tACS effects

To assess whether tACS modulates perceived flicker brightness in a phase-dependent manner, we computed the proportion of brighter ratings of the LED_8Hz_ relative to the LED_12Hz_ for each of the six LED_8Hz_-tACS phase shift conditions. To quantify the degree of tACS phase-dependence, we applied a parametric alignment-based method proposed by Riecke et al. [36] and Zoefel at et. [37]. Herein, the maximum brightness rating condition was aligned to the center bin and the remaining conditions were phase-wrapped. Aligning data to one phase bin naturally skews variability which could produce false positive results if not corrected for. Thus, the average brightness rating of the two bins adjacent to the bin opposite to the center bin (ADJ) was subtracted from the average brightness rating of the two bins adjacent to the center bin (MAX, Fig. 4A). In case of more than one optimal phase bin per participant, we repeatedly computed MAX-ADJ values per optimal data bin alignment and conservatively took the average of MAX-ADJ values. By excluding the center bin and the bin opposite to the center bin, analytical bias that might account for reported tACS effects was prevented [37,38].

In our experimental setup, stable phase entrainment by visual flicker is the key prerequisite for observing phase-specific modulations of flicker-evoked rhythms by tACS [26]. According to this rationale, we examined the dependency of tACS-induced perceptual effects quantified by MAX-ADJ on the degree of SSR-flicker phase locking for each tACS montage and flicker frequency. The absolute correlation coefficient difference between the occipital and periorbital tACS montage was computed per electrode and tested for statistical significance by cluster permutation statistics. To generate the permutation distribution, correlation coefficients were repeatedly computed after pairwise shuffling of MAX-ADJ and SSR-flicker phase locking values between tACS montages per participant across 10,000 permutations. Electrodes were considered significant when the observed correlation difference exceeded 97.5 % of the permutation distribution. For an overall test of the tACS efficacy on the final electrode cluster, we used a linear mixed model to analyze the effect of the fixed factors “tACS montage” (periorbital vs. occipital), “SSR-flicker phase locking at 8 Hz”, “SSR-flicker phase locking at 12 Hz” as well as the interaction effects between SSR-flicker phase locking and tACS montage on brightness modulation. PLVs were averaged across the previously defined cluster electrodes. The subject was set as random effect to account for differences in overall perceptual ratings independent of experimental manipulation. Bonferroni correction was used to adjust for multiple comparisons during post hoc testing. Model fitting was implemented using SPSS Statistics 27 (IBM Corp.).

To investigate the predictability of the optimal timing of tACS application, we computed circular correlation coefficients between the individual SSR-flicker phase delay and the optimal flicker-tACS phase shift leading to greatest brightness ratings. Therefore, we fit one-cycle sine waves to brightness ratings and extracted the optimal phase corresponding to the peak of the sine. Statistical significance of the correlation coefficient per electrode was tested by computing permutation distributions of correlation coefficients between individual SSR phase delays and randomly assigned optimal phase values across 10,000 iterations. The observed circular correlation was considered statistically significant when exceeding the upper 5 % of the permuted distribution.

## Results

### Individualization of physical flicker luminance intensity

In order to account for interindividual differences in flicker frequency-dependent brightness enhancement, we first determined the discrimination threshold of perceptual differences between the LED_8Hz_ and the LED_12Hz_ without tACS. Therefore, we fit pychometric functions to the LED brightness discrimination performance at varying LED luminance ratios per participant (Fig. 2A). The mean luminance ratio between LED_8Hz_ and LED_12Hz_ at the 50 % discrimination threshold was .96 with a mean standard error of .036 (Fig. 2B). Thus, on average participants perceived the LED_8Hz_ as slightly brighter compared to the LED_12Hz_. The psychometric function was estimated on both testing sessions with high retest-reliability, as shown by a significant correlation between the mean luminance ratios within the 35 % and 65 % discrimination thresholds estimated per session (*r* = .84, *p* < .001). Yet, to ensure identical visual stimulation for the two tACS montages, participants were presented with luminance ratios determined on the first day on both testing sessions. Median peak luminance values for the LED_8Hz_ ranged between 78 and 102 cd/m^2^ across brightness discrimination thresholds, while the LED_12Hz_ had a constant peak luminance of 95 cd/m^2^.

**Fig. 2.**
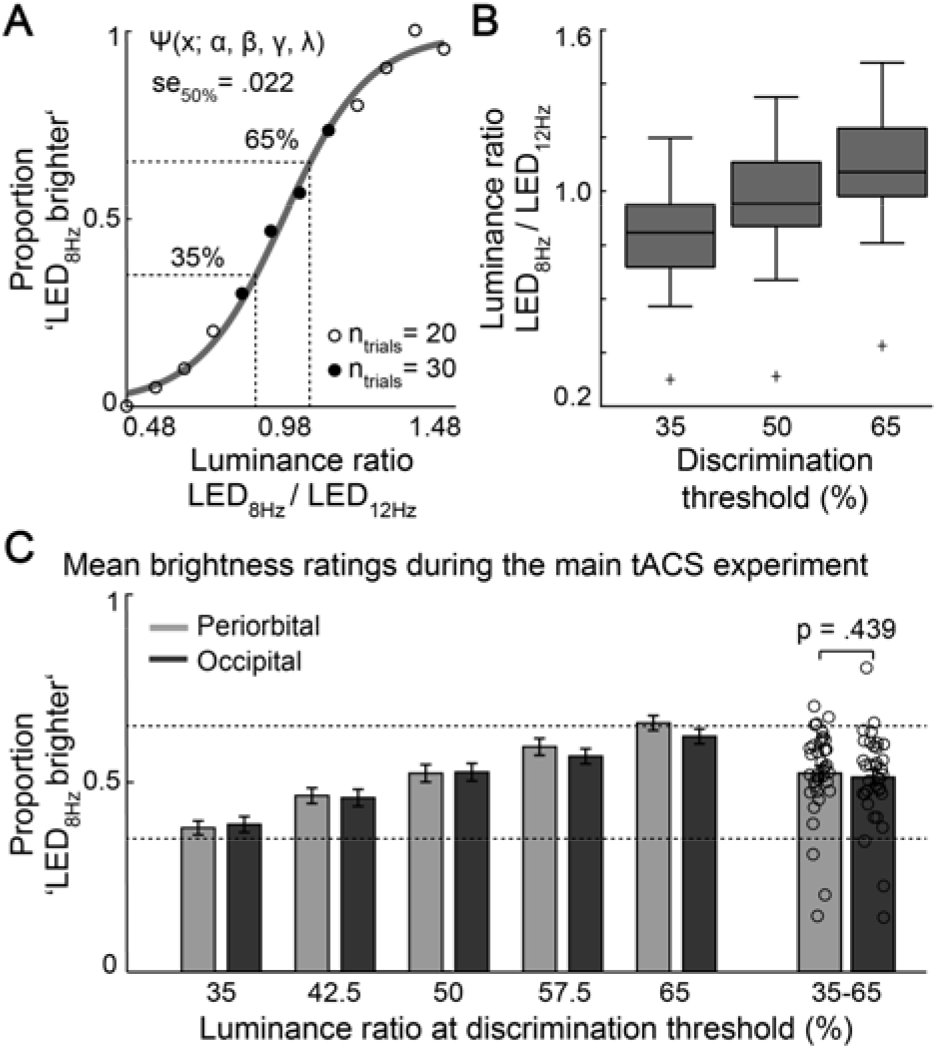
Individualization of physical luminance intensities around flicker brightness discrimination thresholds. (A) Psychometric function for one exemplary participant. Individual luminance ratios between LED_8Hz_ and LED_12Hz_ were determined between the 35 and 65 % brightness discrimination thresholds. (B) Distribution of the estimated luminance ratios between LED_8Hz_ and LED_12Hz_ across participants. (C) Bar diagram shows the proportion of trials on which the LED_8Hz_ was judged as brighter, averaged across LED_8Hz_-tACS phase shifts during the main tACS experiment. As intended, depending on the luminance ratio between LED_8Hz_ and LED_12Hz_, performance ranged between 35 and 65 % brighter judgements. There was no statistical difference between average judgements during the occipital and periorbital tACS testing session.

For the main tACS experiment, trials were presented at five luminance ratios around discrimination threshold, at each of the six tACS-LED_8Hz_ phase shifts. To verify that participants accurately performed the brightness discrimination task, we averaged the proportion of brighter judgements of the LED_8Hz_ across tACS-LED_8Hz_ phase shift conditions, separately for each testing session. As shown in Fig. 2C, the proportion of trials on which the LED_8Hz_ was rated as brighter ranged between 35 % and 65 % of judgements. Without taking flicker-tACS phase shifts into account, there was no significant difference between mean brighter ratings in the occipital and periorbital tACS condition across luminance ratios (*t*(35) = −0.78, *p* = .439, *d_z_* = −.13).

### Interindividual differences in phase stability of visually evoked responses

A basic prerequisite for phase-specific interaction between visual and electric neuromodulation is the phase stability of the targeted flicker-evoked responses. We examined individual sensory entrainment by single flicker stimulation in the right lower visual field with spectral analysis of EEG data that was recorded prior to electrical stimulation. Amplitude spectra during visual stimulation showed peaks at the flicker stimulation frequencies and its harmonics (Fig. 3A). To check for evoked neural phase stability, we computed the degree of SSR-flicker phase alignment for each participant (Fig. 3B). Phase-locking values (PLVs) across all parieto-occipital electrodes were positively correlated between the two testing days (8 Hz: *r* = .61, *p* < .001; 12 Hz: *r* = .59, *p* < .001). Repeated measures ANOVA revealed that flicker phase entrainment was comparable between sessions and flicker stimulation frequencies (frequency: *F*(1,35) = .81, *p* = .373, 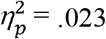; session: *F*(1,35) = .64, *p* = .428, 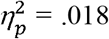; frequency*session: *F*(1,35) = .08, *p* = .782, 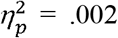). Thus, the two testing sessions did not differ significantly with respect to the detected brain state being targeted by tACS. Topographies of PLVs reflect the spatial specificity of electrophysiological flicker responses, with maximum SSR-flicker phase alignment in the hemisphere contralateral to the flickering stimulus presented in the right visual field (Fig. 3C).

**Fig. 3.**
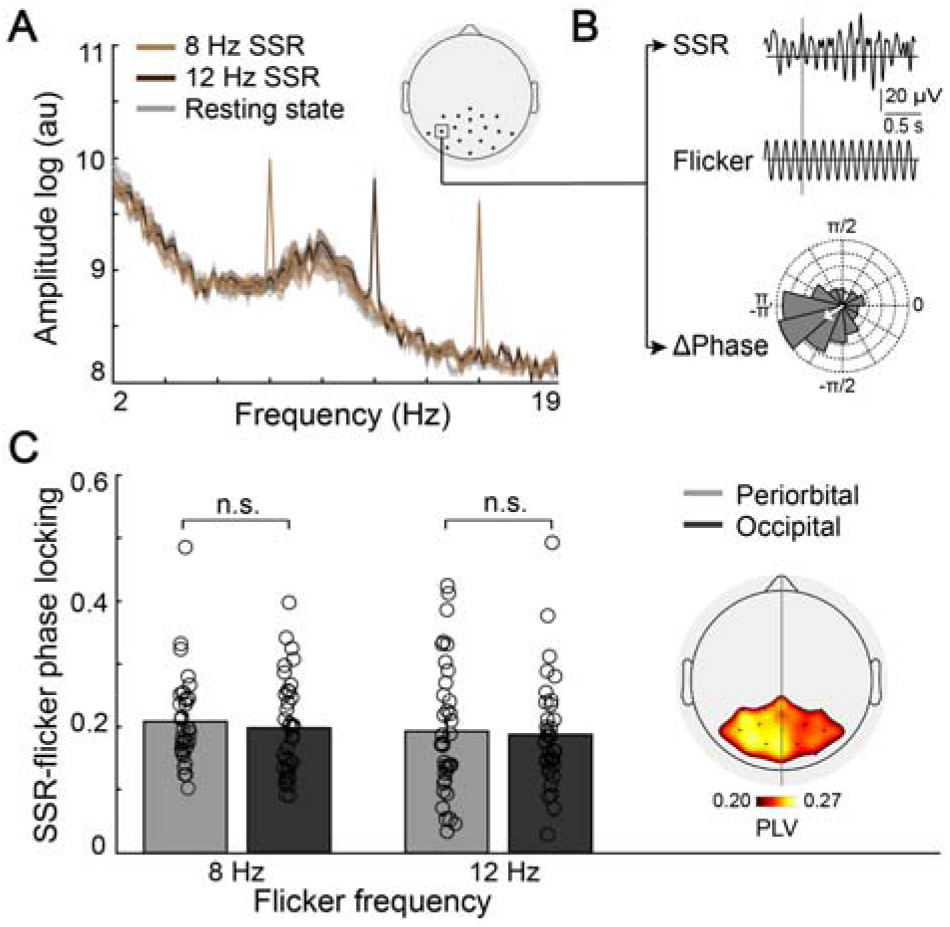
Sensory entrainment by visual flicker. (A) EEG amplitude spectra averaged across participants reveal peaks at the stimulation frequencies of 8 and 12 Hz (and the first harmonic at 16 Hz). Amplitude values were averaged across all recorded parieto-occipital electrodes. (B) Steady-state response (SSR) during 8 Hz flicker stimulation shown exemplary for one participant at electrode PO7. The absolute value of the mean time-dependent differences between SSR and flicker phase reveal the strength of phase alignment by visual flicker. (C) Phase alignment to flicker frequencies averaged across all electrodes did not differ between the occipital and periorbital tACS condition. Insert on the right shows the mean topography of EEG-flicker phase locking values (PLVs) averaged across both flicker frequencies, testing sessions and participants. Phase alignment is highest in visual areas of the left hemisphere contralateral to the flickering stimulus.

### Phase-dependent modulation of flicker brightness perception by occipital tACS

To investigate the degree of perceived flicker brightness modulation by the phase-shift between visual flicker and tACS, we examined the proportion of brighter ratings of the LED_8Hz_ relative to the LED_12Hz_ for each of the six LED_8Hz_-tACS phase shift conditions. tACS phase-dependent perceptual modulation was quantified by a parametric alignment-based method (MAX-ADJ), with values greater than zero reflecting a cyclic modulation of brightness ratings across flicker-tACS phase shifts (Fig. 4A). As stable flicker phase entrainment is the prerequisite for phase-specific flicker-tACS interaction, we investigated the dependency of the tACS effect on SSR-flicker phase locking per flicker frequency. Only for the 8 Hz flicker, permutation statistics revealed an electrode cluster showing a significant difference between tACS montages in the predictability of perceptual modulations by SSR-flicker phase locking (mean cluster correlation and *p*-value: *r* = 0.26, *p* = .009; topography in Fig. 4B, left). The cluster was located in the left hemisphere, contralateral to the flickering stimulus. Fig. 4B shows the relation between tACS-induced brightness modulation and SSR-flicker phase locking averaged across cluster electrodes. For this electrode cluster, the frequency- and montage-specificity of tACS effects was further examined in a linear mixed model analysis. Analysis revealed a significant interaction effect between fixed factors “tACS montage” and “SSR-flicker phase locking at 8 Hz” (*F*(1, 40.89) = 6.19; *p* = .017). All other main and interaction effects were non-significant (see Supplementary Material B, Table B1). The statistically significant interaction term was broken down by conducting separate models for the two “tACS montage” groups on the fixed factor “SSR-flicker phase locking at 8 Hz”. Analysis showed that under occipital electrical stimulation, flicker entrainment strength significantly predicted the degree of perceptual modulation (*F*(1, 34) = 7.59; *p* = .009 < Bonferroni corrected α = .025). However, under periorbital tACS, SSR-flicker phase locking had no predictive value for the tACS efficacy (*F*(1, 34) = 0.81; *p* = .376) (Fig. 4B). Thus, the dependency of the tACS effect on SSR-flicker phase locking was specific for the occipital stimulation montage, the tACS-modulated flicker frequency at 8 Hz and showed a spatially specific cluster-distribution contralateral to the visual flicker.

**Fig. 4.**
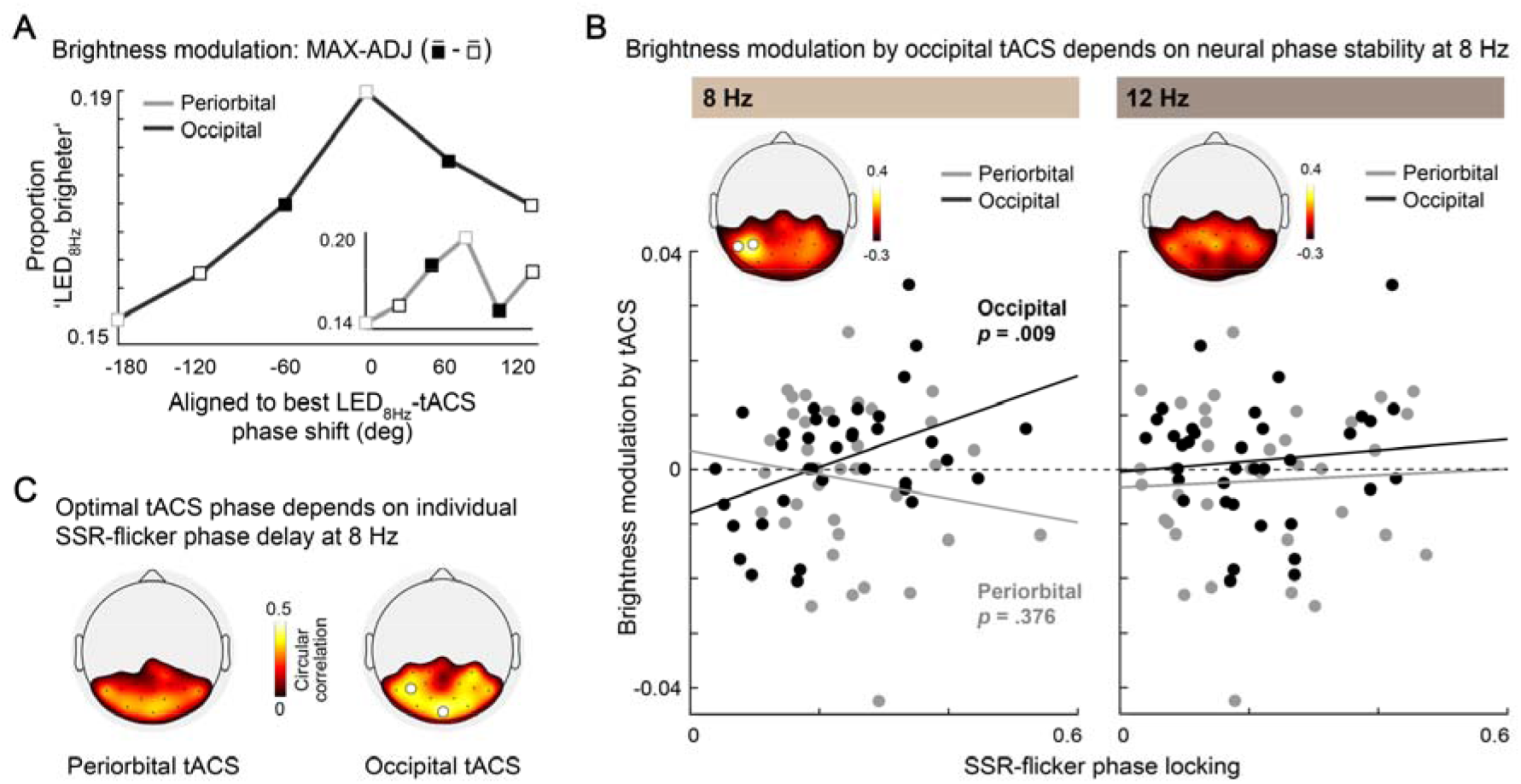
Phase-dependent modulation of flicker brightness perception by same-frequency tACS over the occipital cortex. (A) Proportion of brighter judgements of the LED_8Hz_ compared to LED_12Hz_ for one exemplary participant under occipital tACS. The best LED_8Hz_-tACS phase shift associated with highest brightness ratings was aligned to the center bin (0°). For statistical analysis, phase-dependent modulation was quantified by the MAX-ADJ measure as illustrated. The insert shows data for the periorbital tACS montage. (B) Only for the 8 Hz flicker, an electrode cluster showed a significant difference between tACS montages in the predictability of tACS-induced perceptual effects ( MAX-ADJ) by phase locking of steady-state responses (SSR) to visual flicker. Based on this electrode cluster, linear mixed model analysis revealed a significant interaction between tACS montage and SSR-flicker phase locking at 8 Hz. While brightness modulation under periorbital tACS did not relate to neural phase stability at 8 Hz, occipital tACS effects were dependent on successful flicker phase alignment (*p* = .009 < Bonferroni corrected α = .025). (C) The optimal flicker-tACS phase shift associated with greatest brightness judgements of the LED_8Hz_ was correlated with the individual phase delay between LED_8Hz_ onset and cortical SSR at 8 Hz.

### Cortical phase delay of the SSR predicts the optimal timing of tACS application

While tACS is assumed to influence neuronal excitability with near zero phase lag, cortical SSRs show variable phase lags relative to flicker onset. Accordingly, the optimal tACS-flicker phase shift that enhances brightness perception is expected to vary across subjects. Circular correlation analysis showed that the SSR-flicker phase delay was predictive of the optimal tACS phase, only under occipital tACS (mean electrode correlation and *p*-value: *r* = .31, *p* = .028) (Fig. 4C).

## Discussion

In this multimodal study, we showed that biasing the temporal pattern of cortical excitability by tACS can influence the subjective brightness perception of upcoming rhythmic visual input. Critically, this phase-specific modulatory effect of tACS was only found when electrically stimulating the visual cortex, but not for the retinal control montage. The degree of modulation depended on stable neural phase-locking to the targeted visual flicker and the delay between visual presentation and cortical SSR. Taken together, our data are best explained by a phase-specific cortical interaction between flicker- and tACS-entrained rhythms, causally modulating neural temporal dynamics and related perception.

In this study, the combination of rhythmic visual and electrical stimulation allowed the investigation of tACS effects with control over the phase of the targeted neural rhythm. Previous studies assessed the influence of tACS on ongoing or task-related activity in visual cortex [29,39-44] with only low neural frequency- and phase-consistency of the targeted process, possibly hampering rigorous investigations of neural entrainment effects. In contrast, visual flicker stimulation enables the setting of the oscillatory phase in the visual cortex with high signal-to-noise ratio. Irrespective of whether SSRs reflect entrained intrinsic oscillations or repetitive evoked neural responses [17,45,46], they allow a precise targeting by tACS. In this highly controlled stimulation paradigm, our data show that occipital tACS biases brightness perception in a frequency- and phase-specific manner.

Specifically, perceptual changes were positively correlated with the strength of neuronal phase locking to the tACS-targeted 8 Hz flicker, but not to the 12 Hz reference flicker. This finding supports neural phase stability as the prerequisite for observing phase-specific interactions between SSR and applied electric field [26], and favors an interaction of entrained same-frequency rhythms. Importantly, mean brightness ratings across flicker-tACS phase shifts did not differ between occipital and periorbital tACS, implying a phase-dependent enhancement and reduction of perceived brightness under occipital electrical stimulation. In the same vein, our previous study demonstrated enlarged and suppressed SSR amplitudes depending on the flicker-tACS phase shift [26]. These findings conform with invasive recordings in animals, showing that tACS affects the timing but not the overall rate of spiking activity [23-25]. Taken together, the clear relation between tACS-flicker phase shift and brightness perception strongly points towards an interaction of entrained rhythms in visual cortex that can give rise to SSR amplitude changes and drive perceptual modulations.

The optimal timing of electrical neuromodulation relative to flicker onset leading to highest brightness judgements showed pronounced interindividual variability (see Supplementary Material B, Figure B2). This is in line with previous studies showing that the optimal tACS phase that enhances perception varies uniformly across participants [36,47,48]. Crucially, entrainment of the temporal pattern of neural activity by tACS is expected to facilitate or hamper concurrent visual processing depending on their relative temporal alignment. While tACS can instantaneously affect neural activity, SSRs typically show varying phase delays relative to flicker onset, likely due to interindividual differences in SSR propagation along the visual pathway and anatomical differences [49,50]. In the present data, circular correlation analysis showed a predictive value of the cortical 8 Hz SSR phase delay for the optimal timing of tACS application, exclusively for the occipital tACS montage. As the predictive value of the cortical phase cannot be explained by an interaction between visual input and electric field in the retina, the correlation additionally speaks against a mediation of occipital tACS effects on SSR by retinal co-stimulation. Rather, these data are in support of a neuronal interaction of visually and electrically driven neuromodulation at a cortical level.

By interfering with cortical visual processing via tACS while keeping physical luminance properties constant, our data corroborate potential mechanistic insight into the neural substrate of perception. As tACS has previously been shown to induce shifts in neuronal spike timing [23,25], applied electric fields in our study are expected to have biased the temporal pattern of cortical excitability. These excitability modulations may influence the net synchrony in population firing to upcoming visual input, that generates SSR amplitude changes on a neural population level. Flicker-evoked response amplitudes have been repeatedly shown to correlate with the subjective experience of brightness [9,13,14]. As electrophysiological recordings during the application of tACS are contaminated by electrical artifacts, the direct measurement of SSR amplitude changes during tACS was not possible [51,52]. However, our simulations of neuronal firing patterns to concurrent dual flicker and tACS input (Supplementary Material A) as well as our previous study results [26] demonstrate that this SSR amplitude modulation is feasible, and that it depends on the degree of phase alignment of neural activity to the flicker. The behavioral data observed here reproduce this relation between phase stability of tACS-targeted SSRs and perceptual modulations. Neural populations responding only weakly to visual input might be particularly susceptible to low intensity tACS [53], potentially limiting the positive correlation between SSR-flicker phase locking and tACS effects for large PLVs (see Supplementary Material A and [26]). Thus, extending previous correlative results on the physiological substrate of perception, data suggest a causal relation between brightness perception and the amplitude of rhythmic neural population activity in visual cortex.

A further mechanism that may additionally influence neural responsiveness to visual input builds on pulsed inhibition of evoked responses by tACS-entrained intrinsic alpha oscillations. Besides oscillatory amplitudes, the phase of ongoing alpha rhythms is assumed to exert a cyclic inhibitory influence on neuronal excitability [54–57]. Evidence has been provided for a relation between pre-stimulus alpha phase and subsequent event-related potentials [55,58–60] as well as detection rates of visual stimuli [61,62]. Accordingly, the systematic flicker-tACS phase shifts may have varied the coincidence of luminance peaks with different phases of the tACS-entrained alpha cycle, generating brain-state dependent effects on evoked responses and brightness perception. To experimentally examine these potential mechanisms of tACS, invasive recordings would be necessary that allow the simultaneous assessment of local field potentials and neuronal spiking patterns. Presumably, tACS may exert its effect via multiple ways of action, including a phase-specific modulatory influence on endogenous alpha rhythms as well as on neural populations encoding sensory information.

For tACS applications, it is crucial to consider the role of potential peripheral stimulation effects that might add to transcranial neuromodulation. Innervation of cranial and peripheral nerves in the skin by tACS may induce additional rhythmic activation in the somatosensory cortex. The application of local anesthetics, like EMLA cream used in our study, can effectively reduce somatosensation [24]. Accordingly, participants’ ratings of skin sensation were on average low to moderate and showed no correlation with the strength of perceptual modulation by tACS (Supplementary Material B, Figure B1). Even more important for visual perception research is the possible co-stimulation of the retinae due to current shunting across the scalp [63-65]. To encounter this methodological confound, our active tACS control montage was explicitly tailored to induce subthreshold stimulation of the retinae. Estimated electric field strengths in the eyes under periorbital tACS were comparable to the expected ocular electric field strength under occipital stimulation. As periorbital tACS had no modulatory influence on brightness perception, data strongly suggest that occipital tACS effects were not mediated by subthreshold stimulation of the retinae.

In conclusion, our findings highlight the importance of temporally coordinated activity in visual cortex for subjective brightness perception. We propose that the phase-specific interaction between flicker- and tACS-related activity shaped subjective brightness perception via amplitude modulations of population activity in visual cortex. Our data corroborate the capability of tACS to transmit perceptually relevant temporal information to the human cortex, but also underlines that successful proof for its efficacy is dependent on individual functional properties, emphasizing the necessity for custom-fit stimulation protocols. Transfer of the advantageous methodological approach applied here to other sensory modalities may further help to advance knowledge on the link between cortical sensory processing and perception. Thus, by controlled modulations of brain signals, tACS can be utilized to deepen our understanding of human brain function in basic and clinical science.

## CRediT authorship contribution statement

**Marina Fiene:** Conceptualization, Methodology, Investigation, Software, Formal analysis, Validation, Visualization, Data Curation, Writing - Original draft, Writing - Review & Editing. **Jan-Ole Radecke:** Conceptualization, Methodology, Writing - Review & Editing. **Jonas Misselhorn:** Conceptualization, Methodology, Writing - Review & Editing. **Malte Sengelmann:** Conceptualization, Methodology, Writing - Review & Editing. **Christoph S. Herrmann:** Conceptualization, Methodology, Writing - Review & Editing. **Till R. Schneider:** Conceptualization, Methodology, Writing - Review & Editing, Funding acquisition. **Bettina C. Schwab:** Conceptualization, Methodology, Writing - Review & Editing, Supervision. **Andreas K. Engel:** Conceptualization, Methodology, Writing - Review & Editing, Funding acquisition, Project administration.

## Acknowledgements

This work was supported by the Deutsche Forschungsgemeinschaft (SFB 936/A3 awarded to A.K.E. and T.R.S.; SPP 1665/EN 533/13-1 and SFB TRR 169/B1 awarded to A.K.E.; SPP 1665/SCHN 1511/1-2 awarded to T.R.S.) and by the Studienstiftung des deutschen Volkes (awarded to M.F.). We thank Karin Deazle and Rebecca Burke for assistance in data recording, Florian Pieper for methodological support and Florian Göschl for helpful discussions on the data.

## Supplementary Material A

### Experimental setup

Participants were seated in an electromagnetically shielded, dimly lit room at a distance of 50 cm from the testing apparatus. Two light emitting diodes (LEDs) were placed at 3.4° visual angle in the lower visual field at 6.8° eccentricity from a central fixation dot and a polar angle of 45° from the horizontal meridian (Fig. 1C). Diode LEDs had a diameter of 4 mm and were mounted in color mixing TIR lenses for sharp cut-off light distribution. Lenses were covered by light diffusion paper sheets, yielding a luminous surface with a diameter of 3 cm. The LEDs had a flicker frequency at 8 and 12 Hz. We assessed the nonlinear relation between forward voltage and luminance of each LED by a luminance meter (LS-100, Konica Minolta) and multiplied the resulting function with a pure 8 or 12 Hz sine to generate sinusoidal luminance signals.

### Psychophysical estimation of brightness discrimination thresholds

First, we determined the point of subjective equality between perceived brightness of the 8 and 12 Hz flicker. Based on previous studies showing that subjective brightness perception varies with the temporal modulation frequency [1,2], the point of subjective equality was estimated individually for each participant by applying the method of limits [3]. During eight blocks, the LED_8Hz_ ascended from low to high intensity or descended from high to low intensity, relative to the LED_12Hz_. The 12 Hz flicker was presented on the right side during the first four blocks and on the left side during the last four blocks. Single flicker trials had a duration of 2 sec and were followed by a 1 sec break. Blocks were interrupted by a 3 sec break. Participants responded per button press whenever they perceived brightness of the LED_8Hz_ matching brightness of the LED_12Hz_. Within each block, variable starting and ending luminance intensities were used to avoid expectation errors [3]. The LED_12Hz_ was presented with constant peak luminance of 95 cd/m^2^ (corresponding to 0.4 V peak-to-peak), while the 15 luminance intensities of the LED_8Hz_ ranged between 30 and 160 cd/m^2^ (corresponding to 0.12 – 0.68 V peak-to-peak). Luminance was assessed by a luminance meter (LS-100, Konica Minolta). We averaged the luminance intensities of the LED_8Hz_ selected by participants across 8 blocks to determine the point of subjective equality between perceived brightness at the two flicker frequencies.

Second, we estimated the psychometric function of the proportion of brighter judgements of the LED_8Hz_ relative to the LED_12Hz_ depending on their luminance ratio. 12 evenly distributed luminance ratios, ranging from the priorly estimated point of subjective equality ± 0.5 V, were presented in random order by using the method of constants [3]. 20 trials each were presented for the four highest and lowest stimulus intensities, and 30 trials for the intermediate intensities (Fig. 2A). Single flicker trials had a duration of 2 sec, followed by a break of 1.75 - 2 sec, comparable to the main tACS experiment. After each trial, participants indicated per button press which of the two flickers appeared as brighter (or darker, depending on the assigned condition that was counterbalanced across participants). We fit a logistic function to each participant’s data using a maximum likelihood criterion in the Palamedes toolbox [4]. Guess and lapse rate were fixed to .01 and the luminance ratios determined at [35, 42.5, 50, 57.5, 65] % brightness discrimination thresholds (Fig. 2A).

### Electric field simulation

For electric field simulation, a geometry-adapted hexahedral finite-element head model was computed based on the ICBM-NY “New York Head” [5]. Point electrodes were simulated [6] for the occipital and the periorbital tACS montages, respectively, and electric field simulations were computed with the Simbio toolbox [7]. Electric field strength 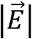 was quantified as the vector length at each location 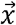 of the electric field 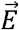 as 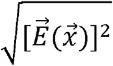. Resulting values in gray or white matter tissue were interpolated on the smoothed cortical surface of the ICBM-NY using a spatial gaussian filter (5 mm width; Fig. 1D, left). Target electric field strength 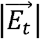 was computed as the average electric field strength in the putative stimulation targets 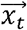 in the left and right visual cortex (Fig. 1D, right). Target locations 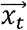 were defined based on retinotopic maps of alpha oscillations evoked by visual stimuli in the lower quadrants of the left and right visual field [8]. To estimate the electric field in the eyes, the surfaces of the vitreous bodies of the eyes were extracted from the segmentation of cerebrospinal fluid within the orbital cavity. Electric field strength 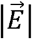 in cerebrospinal fluid was interpolated on these surfaces using a spatial gaussian filter (5 mm width; Fig. 1E, left). Final electric field strength 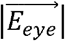 was computed as the 0.95-percentile across all surface nodes of both vitreous bodies, scaled to the individual intensities of the periorbital stimulation and to the common intensity of the occipital stimulation montage (Fig. 1E, right).

### Neural network simulation

To corroborate our behavioral findings, we used a simple neural network model of spiking neurons and simulated the effect of dual visual and electric sinusoidal inputs on SSR amplitudes. Thereby, we aimed to elucidate whether perceptual modulations across flicker-tACS phase shifts would be reasonable at all in case of low neural flicker entrainment. The model was based on Izhikevich [9] and consisted of 1000 neurons [10] with random connections between excitatory (80 %) and inhibitory (20 %) neurons without plasticity of synapses. Each excitatory neuron was connected to 100 random neurons and each inhibitory neuron was connected to 100 excitatory neurons only. Synaptic conduction delays between 1 and 10 ms were assigned to excitatory connections and delays of 1 ms to inhibitory connections. Behavior of each neuron was simulated for 3 sec at a time step of 0.5 ms following the ordinary differential equations:

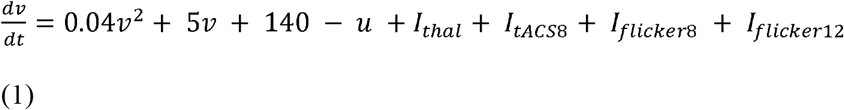

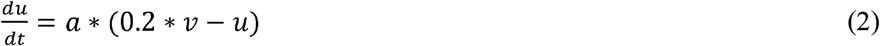

if *v* ≥ 30 *mV*, then *v* = −65,*u* = *u* + *d*

with the neuronal membrane potential *v* and the membrane recovery variable *u*. The time scale and after-spike reset of the recovery variable were set to *a* = .04 and *d* = 8 for excitatory neurons and *a* = .2 and *d* = 2 for inhibitory neurons. Although the model is dimensionless, transmembrane voltage *v* and time *t* can be interpreted in units of mV and ms, respectively. *I_thal_* represents the thalamic input to the neurons, with 10 random neurons receiving an input of 20 every millisecond. The network had mean firing rates of 11 Hz for excitatory and 41 Hz for inhibitory neurons. To simulate the influence of electrical and visual stimulation on neuronal spiking behavior, the following three inputs were added:

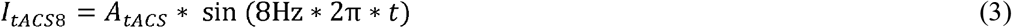

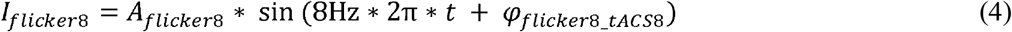

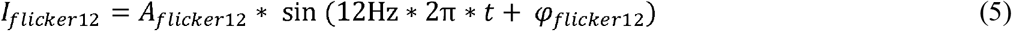

with the amplitudes *A* and the phase shift *φ* between the flicker and tACS signal. *I*_*tACS*8_ was given to all excitatory neurons, while flicker input was assigned to a logarithmically increasing percentage of excitatory neurons from 1 to 100 % to systematically simulate varying levels of flicker-evoked phase synchronization within the network. We quantified the strength of flicker entrainment by simulating neural spiking behavior with *A_tACS_*= 0 and computing the phase locking between the mean transmembrane voltage averaged across the last 2 sec of 800 excitatory neurons and the 8 Hz and 12 Hz flicker sine, respectively. Then, we added *I*_*tACS*8_ with a fixed amplitude ratio of 1:10 relative to the flicker inputs. The phase shift between *I*_*tACS*8_ and *I*_*flicker*8_ was systematically varied (*φ*_*fiicker*8_*tACS*8_: 0°, 60°, 120°, 180°, 240°, 300°). *I*_*flicker*12_ was started at one four randomly chosen phases relative to tACS (*φ*_*fiicker*12_: 0°, 90°, 180°, 270°). To compute resulting SSR amplitudes, we demeaned voltage traces of the last 2 sec epoch per neuron and averaged spectra across all excitatory neurons.

**Fig. S1.**
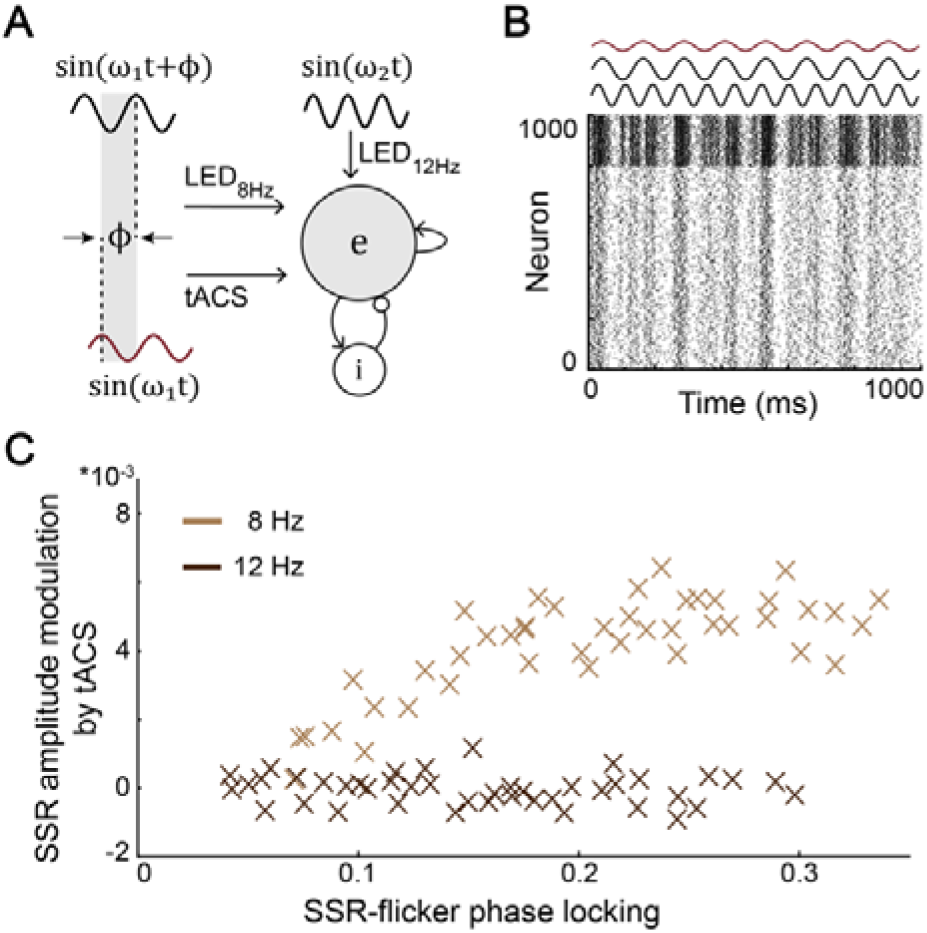
Mechanistic view on the dependency between phase-specific flicker-tACS interference and network synchronization by visual flicker. (A) Architecture of a simple model of spiking neurons, simulating the influence of concurrent tACS and flicker input on neuronal activity. 8Hz flicker and tACS inputs were added with systematic phase shift φ. (B) Neuronal firing patterns are shown for 800 excitatory and 200 inhibitory neurons. (C) Simulations suggests that a modulation of neural steady-state response (SSR) amplitudes at 8 Hz is only feasible under stable SSR-flicker phase locking. In support of behavioral results, simulated data suggest a phase-specific interference of visual and electric inputs.

SSR amplitudes were defined as the ratio between the amplitude at the stimulation frequency and mean amplitude at the two neighboring frequencies. SSR amplitudes and phase locking values were averaged across 90 simulations. Modulation of SSR amplitudes dependent on the phase-shift between visual and electric inputs was then quantified by the MAX-AD J measure (Fig. S1).

Converging with behavioral data, our network simulation reveals a dependency of the tACS efficacy, here quantified by the MAX-ADJ value of simulated population level SSR amplitudes, on neural phase stabilty of the tACS-targeted 8 Hz rhythm (Fig. S1 C). SSR-flicker phase locking at the reference frequency of 12 Hz shows no relation to SSR amplitude modulation by tACS. This supports an interaction of same-frequency entrained rhythms. Thereby, network model simulations suggest that the systematic phase-shift between flicker and tACS can only modulate SSR amplitudes and, thus, possibly also related perception, if flicker-driven neural activity depicts a certain level of phase stability over time (see also [11]).

## Supplementary Material B

**Figure B1.**
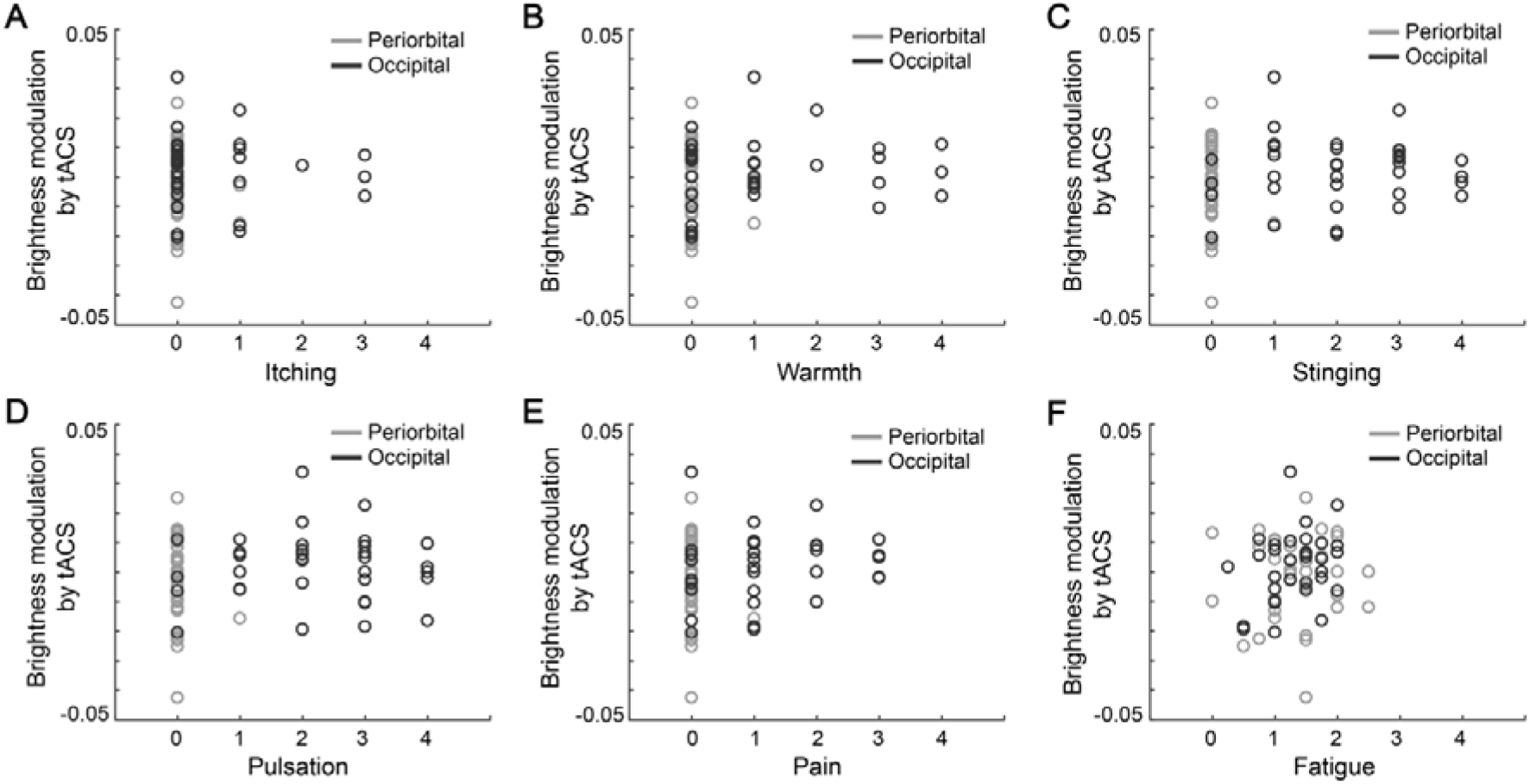
Side effect characteristics during electrical stimulation. At the end of each testing session, participants rated the maximum perceived intensity of skin sensations (itching, warmth, stinging, pulsation), pain and fatigue as either “absent”, “light”, “moderate”, “pronounced” or “strong” (coded from 0 to 4). Fatigue values were averaged across four ratings during the course of each testing session. The relation between the strength of tACS-induced brightness modulation and side effects during periorbital and occipital tACS was tested by Pearson correlation coefficients. None of the correlation coefficients were statistically significant (p > .05).

**Figure B2.**
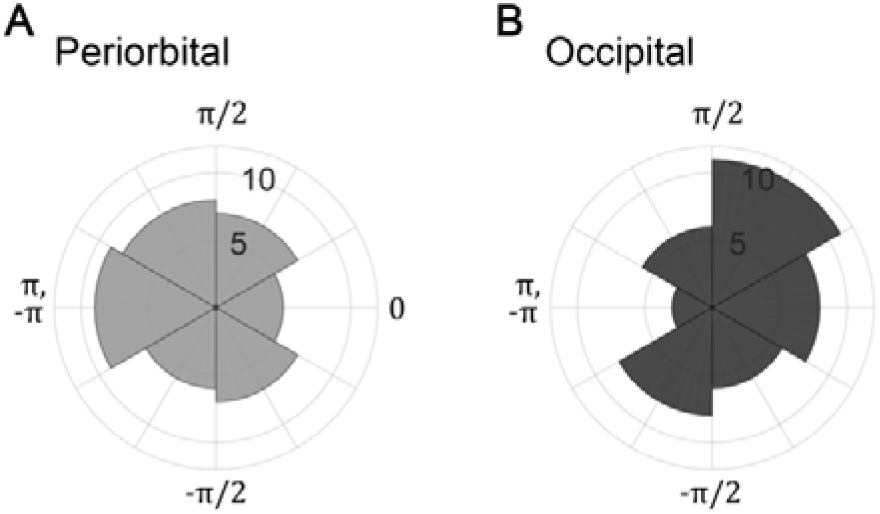
Distribution of optimal tACS-LED_8Hz_ phase shifts leading to highest brightness perception across participants. Neither tACS condition revealed evidence for a nonuniform distribution of optimal phase shifts by Rayleigh tests for nonuniformity. (A) Periorbital tACS: *z*(35) = 0.45, *p* = .639. (B) Occipital tACS: *z*(35) = 1.17, *p* = .313.

**Table B1.**
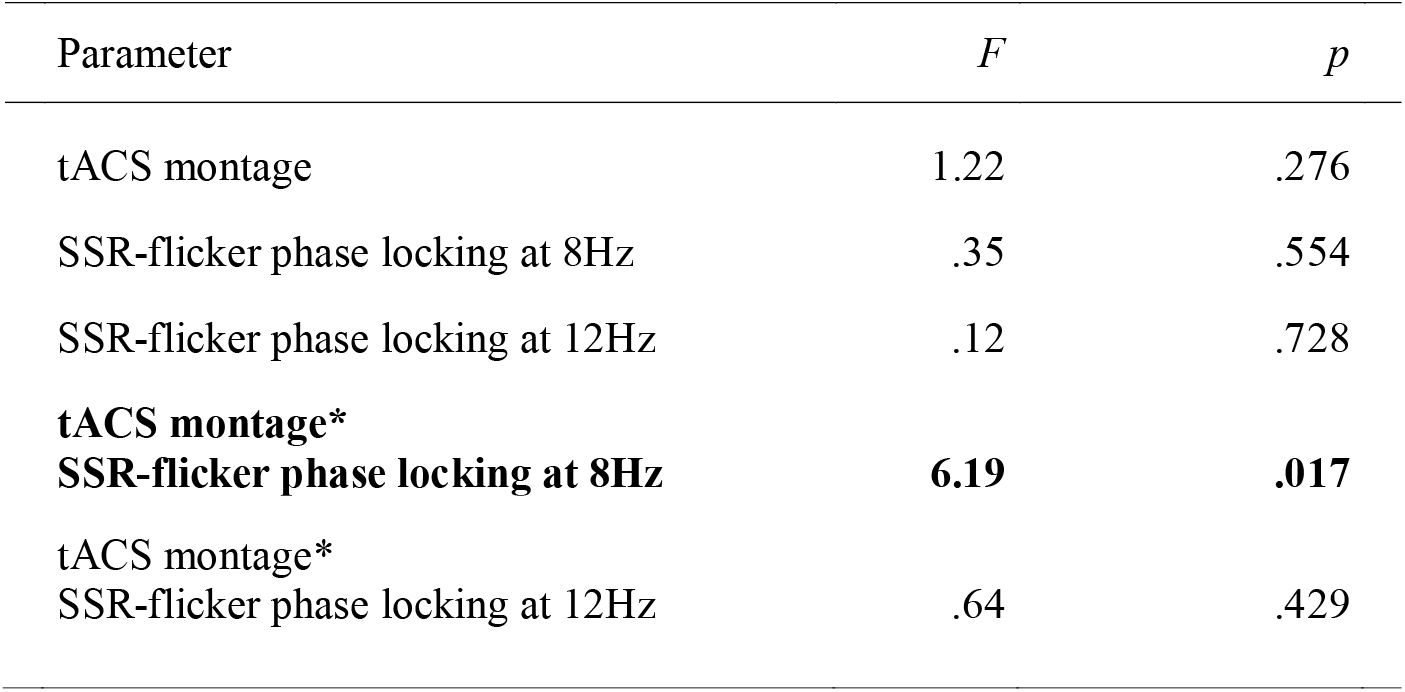
Mixed model analysis results. To investigate the frequency- and montage-specificity of the tACS effect on brightness perception, linear mixed model analysis was computed. “tACS montage” (periorbital vs. occipital), “SSR-flicker phase locking at 8 Hz”, “SSR-flicker phase locking at 12 Hz” as well as the interaction terms between tACS montage and SSR-flicker phase locking were included as fixed factors in the model. Shown are the test statistics for all main and interaction terms. tACS: Transcranial alternating current stimulation; SSR: Steady-state response.

**Table B2.**
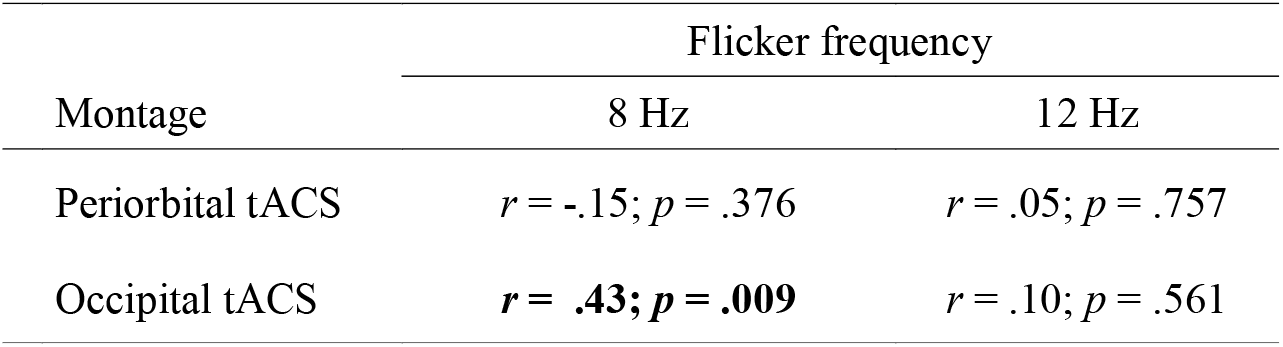
Montage- and frequency-specificity of the tACS effect. Complementary to post hoc linear mixed model results, the table shows Pearson correlation coefficients between the strength of flicker brightness modulation (MAX-ADJ) and phase-locking of steady-state responses (SSR) to visual flicker. The *p*-values are uncorrected for multiple testing. tACS: Transcranial alternating current stimulation.

